# Early-acting inbreeding depression can evolve as an inbreeding avoidance mechanism

**DOI:** 10.1101/2023.03.20.533524

**Authors:** Yaniv Brandvain, Lia Thomson, Tanja Pyhäjärvi

## Abstract

Despite the potential for mechanical, developmental and/or chemical mechanisms to prevent self-fertilization, incidental self-fertilization is inevitable in many predominantly outcrossing species. In such cases, inbreeding can compromise individual fitness. Unquestionably, much of this inbreeding depression is maladaptive. However, we show that when reproductive compensation allows for the replacement of inviable embryos lost early in development, selection can favor deleterious recessive variants that induce “self-sacrificial” death of inbred embryos. Our theoretical results provide numerous testable predictions which could challenge the assumption that inbreeding depression is always maladaptive. Our work is applicable any species that cannot fully avoid inbreeding, exhibits substantial inbreeding depression, and has the potential to compensate embryos lost early in development. In addition to its general applicability our theory suggests that self-sacrificial variants might be responsible for the remarkably low realized selfing rates of gymnosperms with high primary selfing rates, as gymnosperms exhibit strong inbreeding depression, have effective reproductive compensation mechanisms, and cannot evolve chemical self-incompatibility.

## Introduction

Inbreeding depression caused by both rare, recessive, and highly deleterious mutations (1, 2), and by deviations from phenotypic optima under multi-locus stabilizing selection (3–6), is an inevitable consequence of habitual outcrossing. As such, inbreeding avoidance mechanisms, including chemically mediated self-incompatibility (SI) systems, are common across the tree of life (7, 8). The mechanisms and timing of SI are diverse (9), but the best-characterized cases involve an interaction between the diploid maternal genotype and the pollen’s haploid (gametophytic SI) or diploid (sporophytic SI) genotype before fertilization.

Late Acting SI – in which self-pollen can effectively germinate, grow down the stigma, and in some cases can enter and fertilize the ovule before ovular development arrests (10–13) – is less well understood (14). In the most delayed form of Late Acting SI, “Post-zygotic” SI occurs after fertilization (15). Claims of Late Acting SI, and post-zygotic SI in particular, have attracted much attention and criticism both because this mechanism of incompatibility is unusual (14), and because it is difficult to distinguish late-acting SI from early-acting inbreeding depression (9, 16). Traditionally, the key distinction between these mechanisms is that SI is “*a mechanism operated by the maternal parent plant and controlled by the genotypes of that plant and its pollen donor*” and that inbreeding depression is “*a process acting in the progeny zygote, determined by its own genotype*” (16). Substantial effort has gone into distinguishing between early-acting inbreeding depression and late-acting SI (reviewed in 14, 17), with evidence supporting late-acting SI in some taxa (e.g. 18–21) and early inbreeding depression (e.g. 22, 23) in others.

We argue that with high reproductive compensation (i.e. the exchange of dead embryos for potential viable embryos), the distinction between early-acting inbreeding depression and late-acting SI is not necessarily evolutionarily relevant because early-acting inbreeding can adaptively evolve as a mechanism to minimize reproductive investment in low fitness selfed seed. Reproductive compensation is likely common across taxa in which offspring receive parental resources after fertilization, or even when offspring compete among one another for a local resource. The large levels of maternal investment in humans (e.g. 24, 25) and other mammals (e.g. 26) has led to a substantial body of research on reproductive compensation and its consequences in animals. Reproductive compensation is also likely common among angiosperms (27), as most species produce fewer progeny than zygotes (28, 29), a disparity that is elevated in habitually outcrossing species (29). Additionally, reproductive compensation is likely built in to the system of “simple polyembryony” found in gymnosperms, as only one of numerous genetically distinct embryos matures within a seed. As such, in the many taxa with an opportunity for reproductive compensation, an early-expressed recessive load allows mothers to trade in inbred for outbred embryos and can essentially function like an SI system.

Homozygosity for rare recessive alleles serves as a reliable indicator of inbreeding – much like the similarity of pollen and style S-locus genotype serves as a reliable indicator of self vs. non-self pollen. As such, we further argue that all forms of SI can be considered as a special case of this sort of altruistic act as rejected pollen are sacrificing their ability to sire potentially viable seed. Because in many cases early-acting inbreeding depression will reflect the standard expectation of a balance of selection against deleterious alleles and their mutational input (i.e. traditional “mutation-selection-balance”), rather than our novel hypothesis that selection directly favors such alleles, we provide numerous predictions which allow researchers to evaluate whether an early-expressed recessive load reflects selection for, or imperfect selection against it. Since our model makes very few critical assumptions – namely that homozygosity for a rare allele signals recent inbreeding, that inbreeding depression exists, and that early mortality can be compensated, we suggest that many taxa may express an altruistic recessive load.

## Methods

### Overview

We model reproductive compensation as follows – each seed contains a primary and backup embryo, allowing an inviable ‘primary’ embryo to be replaced by another, potentially viable embryo (Figure 1A). Although this model assumes exactly two embryos per seed, our results do not depend on this model of reproductive compensation – in Figure 2 and the Supplementary Material, we show that analytical results from this “backup embryo” model are nearly identical to those obtained by using Charlesworth’s (26), model of reproductive compensation, in which maternal fitness equals 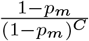, where *p*_*m*_ is the probability that an embryo will not survive and *C* is the extent of reproductive compensation (Described in more details in the Supplementary Text). All code and data associated with this manuscript are available through an associated zenodo repository (30).

**Fig. 1.**
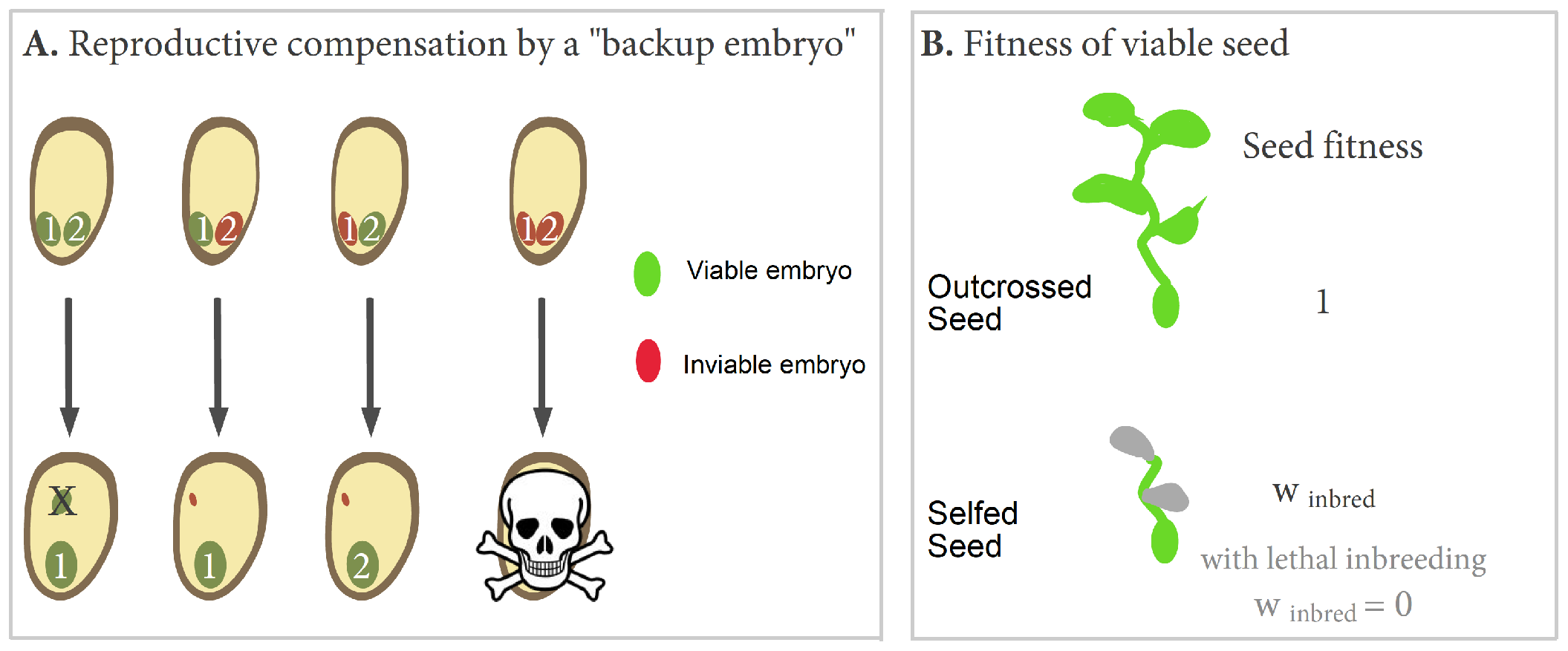
Stages of selection in the “backup embryo” model. Figure 1A depicts our model of reproductive compensation. Here the backup embryo (embryo 2) is eliminated if both embryos are viable (noted in green), dies if inviable (noted in red), and replaces the primary embryo (embryo 1) if the primary embryo is inviable. If both embryos are inviable there is no reproductive compensation. Figure1 B Shows the second stage of selection in which the fitness of viable outcrossed seeds is one, and the fitness of viable selfed seed is *w*_inbred_, which equals 0 in the case of lethal inbreeding.

**Fig. 2.**
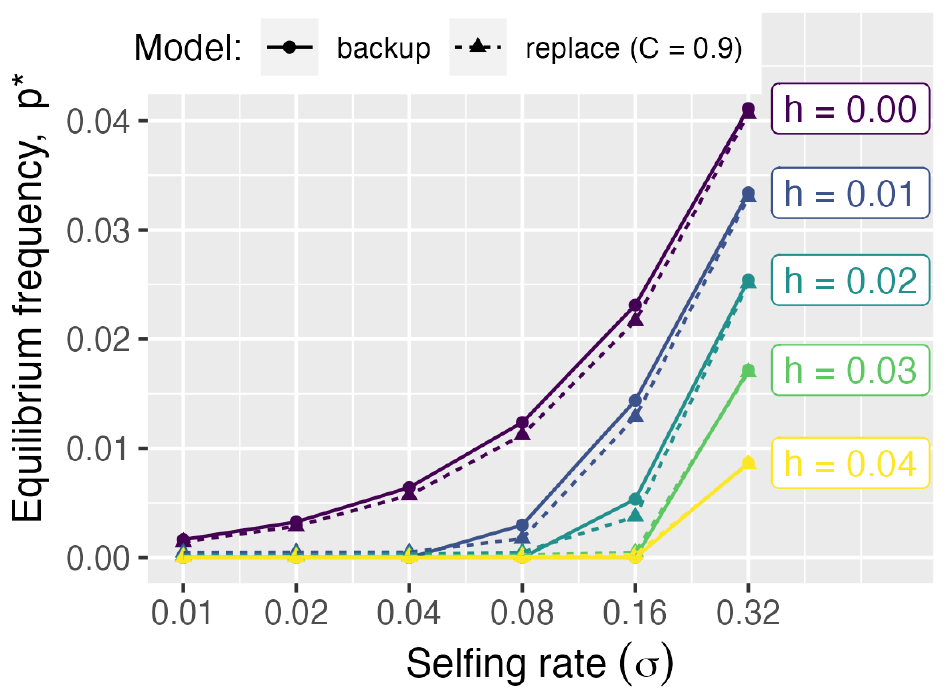
The equilibrium frequency, *p*^*∗*^, of a self-sacrificial (partially) recessive allele for which lost embryos can be compensated, under a variety of selfing rates and dominance coefficients. Results are presented from our “backup embryo” model are presented with circles connected by solid lines). These results are nearly identical to those from Charlesworth’s parameterization of reproductive compensation (triangles connected by dotted lines). in which maternal fitness equals 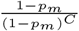, where *p*_*m*_ is the probability that an embryo will not survive and *C* is the extent of reproductive compensation. This figure displays results for *C* = 0.9 and with *s* = 0.6. The difference between results with our and Charlesworth’s parameterization of reproductive compensation widens as *C* diverges from 0.9, with *p*^*∗*^ decreasing as *C* decreases (Figure S4). In this figure, we assume that the allele is lethal when homozygous, however, results are remarkably insensitive to *s* (Figure S5).

Selection occurs at two distinct phases of the lifecycle. Selection first occurs within maternal families based on the recessive allele(s) underlying early embryo viability (i.e. between viable and inviable embryos in Figure 1A), and among maternal families based on number of fully developed seeds per mother (e.g. mothers with inviable primary and backup embryos will experience a reduction in fitness – Figure 1A). Next (after embryos can be compensated) selection occurs based on inbreeding status and the extent of inbreeding depression (Figure 1B). We assume that inbreeding depression is expressed only after seeds complete development (initially all embryos survive to become full seeds), that it is not purged at the population’s mutation and selfing rates. This evolutionary stable load can occur when deleterious recessive alleles accumulate more rapidly than selection eliminates them (31), or when inbreeding leads to predictable deviations away from phenotypic optima in populations experiencing multi-locus stabilizing selection (4–6). Despite the ample evidence for early expressed inbreeding depression in nature, our model assumes its absence because (a) if there is more early inbreeding depression than our model predicts that a ‘self-sacrificial allele’ cannot invade, and if there is less, such an allele can invade with its equilibrium frequency lowered by the amount of pre-existing early inbreeding depression, and (b) Empirical cases of early acting inbreeding depression in the many taxa with a possibility for reproductive compensation could reflect the processes presented in our model.

First, to evaluate whether early expressed deleterious recessive alleles can be favored by natural selection, we analyze deterministic single-locus models without mutation or drift. Second, to evaluate the extent to which the adaptive altruistic load discovered in our single locus deterministic models can contribute meaningfully to the early expressed recessive load in a finite population experiencing recurrent mutation and genetic drift, we conduct stochastic, genome-scale simulations.

### Deterministic Single-Locus Models Without Mutation

We initially assume that all primary embryos survive to become a seed, and that following seed development, seeds die if they are inbred and survive otherwise (we later relax the assumption that inbreeding is lethal). We then introduce a rare, highly recessive (with dominance coefficient, *h*) allele, *A*, which decreases embryo survival by *s*. Embryo survival is then converted into the expected maternal fitness following the model of reproductive compensation employed (Figure 1A,B)

We then explore the invasion of this allele under differing selection and dominance coefficients, and find the equilibrium frequency of such an allele in the absence of mutation. Throughout, we assume a population of infinite size in which mothers which incidentally self-fertilize (with probability, *σ*) or randomly mate (with probability 1 *− σ*). Details of our analytical setup are provided in the appendix.

### Stochastic Genome-Wide Models With Mutation

#### Simulation framework

We developed genome-scale, individual-based stochastic simulations to test whether (and to what extent) the adaptive altruistic load discovered in our analytical model could contribute to the early expressed load in a finite population experiencing recurrent mutation and genetic drift. Specifically, we ran non-Wright-Fisher forward simulations in SLiM 3.0 (32) in which early-acting recessive deleterious mutations recurrently arise (at rate *µ*) at any location in the genome. We followed these simulations until populations achieved mutation-selection balance. We specify a genome of one hundred thousand sites, such that e.g. a mutation rate of 0.5 *×* 10^*−*6^ corresponds to an expectation of one mutation per diploid genome per generation. These sites are uniformly spread over ten chromosomes, each of which is one Morgan in length with a uniform recombination map. We present results in the main text for a population of size ten thousand, and explore how changing this size impacts results in the supplementary material.

We explore results for a variety of selfing rates, and for modest (*µ* = 5 *×* 10^*−*8^) high (*µ* = 5 *×* 10^*−*6^) mutation rates, and for lethal ((*w*_inbred_ initial = 0.00) and sub-lethal ((*w*_inbred_ initial = 0.25) inbreeding depression. Because deterministic results from our “Backup embryo” model are nearly identical to results from those obtained using Charlesworth’s (26) parameterization of reproductive compensation, we limit our computationally intensive whole genome simulations to our primary model of reproductive compensation.

#### Evolving an altruistic load

Each generation, 10,000 survivors were randomly sampled with replacement to act as mothers of two half-sib progeny. Each embryo was inbred (i.e. the product of selfing) with probability *σ*, the product of outcrossing (i.e. outbred) with probability 1 *− σ*. Next, early-acting deleterious mutations were expressed and embryos experienced viability selection based on their embryonic load. Finally, if both siblings survived, one was randomly considered “primary” and became a seed. The fitness of this seed was independent of its genotype and depended exclusively on its inbreeding status.

#### A control

As a control, we developed a second model where the early expressed load was not favored by natural selection. Again, two maternal siblings were produced from 10,000 mothers (sampled with replacement) every generation and embryo fitness depended on early-acting deleterious mutations. Unlike the first model, replacement did not occur between siblings; instead occurring between random embryo pairs. If both embryos in the pair survived, one was randomly considered “primary” and became a seed, this removes the opportunity for altruistic selection to favor an early-expressed genetic load. We finally instituted viability selection on surviving embryos as described above.

## Results

### Selection can favor recessive deleterious alleles

#### Invasion of a self-sacrificial mutation

To evaluate invasion conditions in the “Backup embryo” model, we assume the *A* allele is so rare that *AA* homozygotes can be ignored, so we consider sibling pairs from *aa* or *aA* mothers. Outbred ovules are pollinated by *A* or *a* bearing pollen, with probabilities *p*_*A*_ and 1 *− p*_*A*_, respectively.

During the invasion of the recessive *A* allele, (i.e. lim *p*_*A*_ → 0) *aa* mothers produce *aa* offspring by selfing with probability *σ* and produce *aa* and *Aa* offspring by outcrossing with probabilities (1 *− p*_*A*_)(1 *− σ*), and (*p*_*A*_)(1 *− σ*), respectively. The probability that the primary embryo in an *aa* maternal family is replaced by a backup embryo, *ψ*_*aa*_, equals the product of the probability that an embryo is fertilized by *A* pollen and dies in early development i.e. *hsp*_*A*_ (1 *− σ*) (See our mathematical appendix for details).

At invasion, *Aa* mothers produce *aa, Aa*, and *AA* offspring by selfing with probability 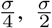 and 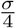, respectively, and produce *aa, Aa*, and *AA* offspring by outcrossing with prob-abilities 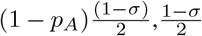, and 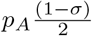, respectively. The probability that the primary embryo in an *Aa* maternal family is replaced by a backup embryo is *ψ*_*Aa*_, and equals 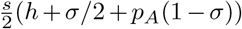.

At invasion, the population mean fitness is 1 *− σ* (as we are assuming lethal inbreeding) and the mean fitness of rare *Aa* maternal families, 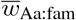 is

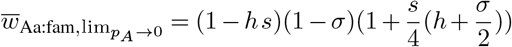

Assuming full recessivity (i.e. *h* = 0), 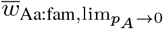 is 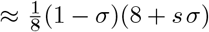, which always exceeds mean population fitness a t invasion, and therefore a recessive *A* is never disfavored when rare. The expected increase in frequency of a rare recessive allele that acts in this manner is 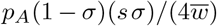. Our results do not depend on the assumption of lethal inbreeding depression – – assuming that all outbred seed survive, and that the load cannot be purged, a recessive allele that decreases embryo fitness invades when the probability that an inbred offspring survives is less than 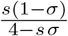.

Similarly, complete recessivity is not required for the deterministic spread of this self-sacrificial allele. With lethal inbreeding, a partially recessive self-sacrificial allele is favored when the primary selfing rate exceeds a critical selfing level, which equals

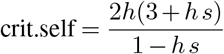

Figure S1 shows that at first approximation, the critical selfing rate required for the invasion of a self-sacrificial allele is linear with its recessivity (*h*), such that it will spread when its selfing rate, *σ*, exceeds 6*h*.

Invasion criteria for a model with Charlesworth’s parameterization of reproductive compensation are nearly identical to those obtained from our backup embryo model (Figures S2 and S3). Specifically, when *s* and *h* are small, invasion criteria from the backup model resembles cases in which reproductive compensation is very effective (between *C* = 0.90 and *C* = 0.99), while the backup model more closely resembles less effective reproductive compensation as the dominance coefficient is larger.

##### Equilibrium frequency of a self-sacrificial mutation

As the self-sacrificial allele increases in frequency, it begins to be exposed commonly in the allozygous, rather than the autozygous, state. As such, as *A* increases in frequency, it becomes a less reliable indicator of inbreeding. To demonstrate this phenomenon, we apply Bayes’ theorem to show that the probability that an embryo was inbred, given that it is homozygous for *A* equals 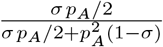 under the assumption of complete recessivity and lethality of the self-sacrificial allele. This results in frequency dependent selection in which the self-sacrificial allele is favored when rare but disfavored when common.

Because our model contains both family structure and departures from Hardy-Weinberg equilibrium, analytically deriving stable intermediate frequency, 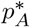, is quite challenging. Rather than deriving 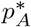, we iterate the genotype frequency recursion equations until genotype frequencies are stable. In both models, *A*’s equilibrium frequency is highest when it is highly recessive and in a frequently self-fertilizing population (Figures 2 and S4), and its equilibrium frequency is lower as its dominance increases or the population’s selfing rate decreases. The selection coefficient, *s* has a nearly no effect on this equilibrium frequency (Fig S5).

#### A genome-wide altruistic load

Is this deterministic selective benefit of self-sacrifice strong enough to overcome the sampling process of random genetic drift? If so, when will the benefit of replacing oneself with a viable half sibling leave an observable pattern on the genetic load, such that it can be separated from the standard process of mutation selection balance? What should the frequency spectrum of such self-sacrificial alleles look like – should it be marked by an excess of relatively rare or relatively common variants? Answers to such questions – which we derive by genome-wide simulation – are essential for our ability to test the adaptive load hypothesis in natural populations.

Figure 3 compares the mean frequency of early-expressed deleterious recessive alleles (*h* = 0) per individual across three selfing rates (*σ* = 0.1, *σ* = 0.2, and *σ* = 0.3, x-axis), two mutation rates (*µ* = 5 *×* 10*−*8, or *µ* = 5 *×* 10*−*6, rows), three selection coefficients (*s* = 0.2, *s* = 0.6, and *s* = 1.0, columns), and two levels of inbreeding depression (*w*_inbred_ initial = 0.00 Fig. 3A or (*w*_inbred_ initial = 0.25, Fig. 3B). Although we present results only for mean allele frequency, we find qualitatively similar results for different summaries of the mutational load (e.g. the mean number of mutations per individual, Fig S6). We find that the mean mutation frequency only weakly depends on the selection coefficient (Fig. 3), a result also found in our single-locus analytical model (Fig S5). The efficacy of selection for an altrusitic load increases with population size – most notably the number of deleterious recessive variant per individual and the mean frequency of such variants is comparable between our model and the control when populations are small (e.g. *N* = 100 or *N* = 500) but these values become more distinct as the population size increases (Figures S7 and S8).

**Fig. 3:**
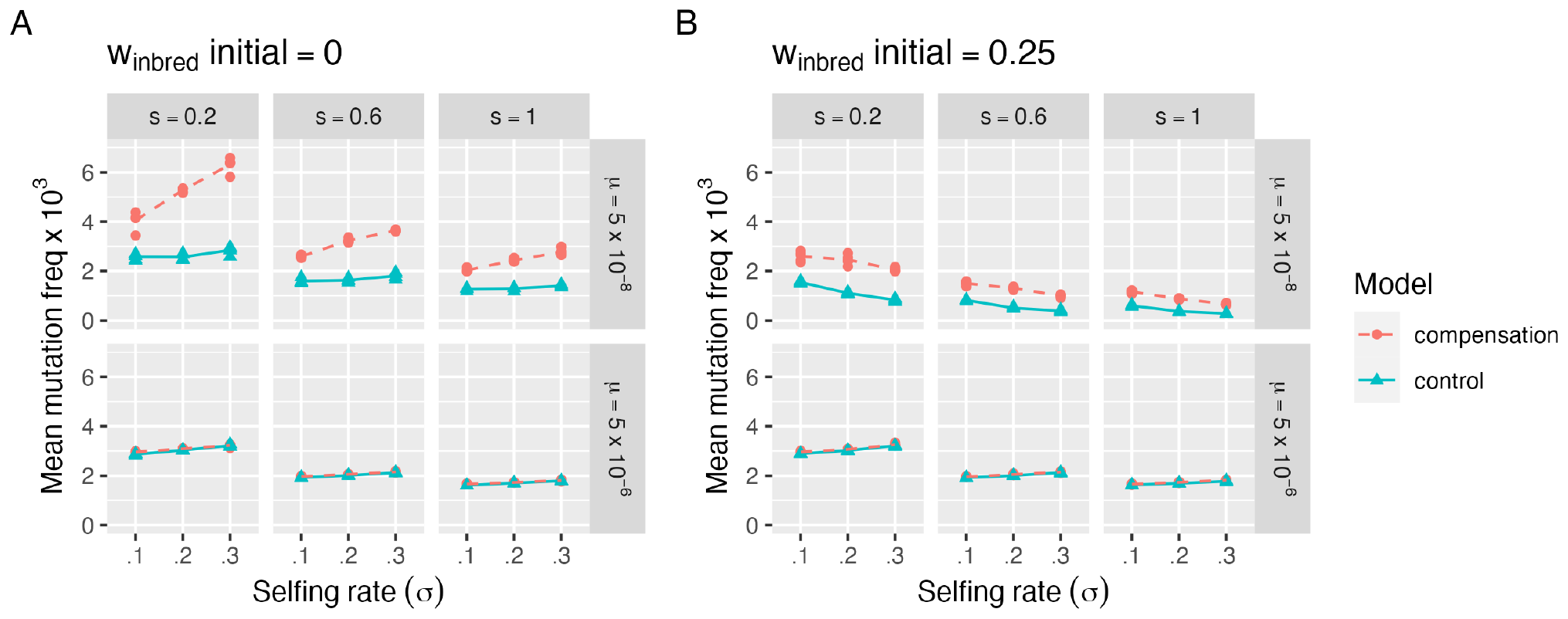
The mean frequency of early-acting, recessive alleles, when inbred embryos have zero or low (*w* = 0.25) fitness before such alleles are introduced (panels A and B, respectively). We present results of a stochastic genome-wide SLiMulation with *N* = 10, 000 over a combination of selfing rates (*σ*), mutation rates (*µ*), and selection (*s*) coefficients. Red dotted lines display results for cases in which primary embryos are replaced by siblings (our model), while blue solid lines display cases in which primary embryos are replaced by a non-sibling (a control). Simulations were run in a population of size 10000.

With lethal inbreeding depression and low mutation rates, there is a clear elevation in the frequency of early-expressed deleterious recessive mutations when reproductive compensation occurs in comparison to when it does not (Top panel of Fig. 3A), consistent with expectations from our analytical theory. This pattern intensifies as the selfing rate increases, presumably because a higher selfing rate increases the necessity of removing selfed embryos. These patterns disappear as the mutation rate increases (Bottom panel of Figure 3A), presumably because the mutational input outpaces what is required to ensure the death of selfed embryos. As such, selection acts to remove low fitness allozygotes (i.e. homozygotes formed by outcrossing). This explanation is consistent with the increased number of such mutations per individual with this higher mutation rate (Fig S6) and their lower frequency when segregating.

With sublethal inbreeding depression (*w*_inbred_ initial = 0.25, 3B), we find a similar elevation in the frequency of these recessive alleles with reproductive compensation as compared to controls. However, in this case, the frequency decreases, rather than increases with the selfing rate. This difference in the relationship between the selfing rate and the frequency of a self-sacrificial recessive allele with lethal versus sublethal inbreeding likely reflects the fact that with sublethal inbreeding depression there is a cost to having two inviable selfed embryos. As such, this result may be idiosyncratic to our model of reproductive compensation.

## Discussion

The opportunity for reproductive compensation is likely pervasive across the tree of life. Egrets lay two eggs per season with only one destined to survive (33). Humans can make up for embryo mortality by increasing reproductive effort (34, but see (35)). Birds tend to lay more eggs during plagues of their rodent predators, and ultimately clutch size in many birds is determined by how many young can be fed, not how many can be laid (36). Similarly, angiosperms produce many more ovules than seeds (37). The evolutionary consequences of reproductive compensation have received much attention. For example, Hurst (38) recently argued that reproductive compensation provides selfish centromeres novel opportunities to functionally distort meiosis. In Hurst’s model, rather than directly driving during meiois, selfish centromeres inducing nondisjunction to sabotage embryos destined no to inherit this centromere, and if such lost embryos can be compensated this sabotage gives the selfish centromere an additional opportunity to solely end up in the egg, rather than the polar body. Additionally many researchers have followed up on the suggestion R.A. Fisher made to Race (39) that reproductive compensation in humans allows for evolutionary stability of polymorphism at the underdominant Rh locus (40, 41). More generally studies in plants (27, 42, 43) and mammals (24, 44) have shown that reproductive compensation results in an increase in the early-expressed recessive load. Our study shows that reproductive compensation not only relaxes selection against deleterious recessives, but actually favors an early-expressed recessive load when inbreeding is unavoidable and inbreeding depression is severe.

Our finding that selection can favor an early expressed recessive load makes sense of Sakai’s (45, 46) observation that reproductive compensation plus both early and late acting deleterious recessive mutations allowed for both mutational types to segregate at higher frequencies than they would on their own. We interpret this “two-stage effect” as an increase in the early expressed load favored by the mechanisms proposed above, followed by the relaxation of selection against late expressed recessive load as the loss of embryos formed by self-fertilization and biparental prevents recessive alleles expressed later in life from being exposed when rare.

We focused on cases of unavoidable self-fertilization in plants. However the logic of our model is more broadly applicable. Although quantitative results will differ based on details of the systems of mating, we hypothesize that an adaptive early-expressed recessive load could evolve in plants and animals with the potential for reproductive compensation and in which dioecy or self-incompatibility cannot eliminate incidental biparental inbreeding and subsequent inbreeding depression (e.g. 47–50).

### Precedent for the adaptive evolution of self-sacrificial alleles

Ours is not the first study to examine the possibility of the evolution of self-sacrificial alleles. For example, cytoplasmic male killers can adaptively spread in maternally transmitted organelles in taxa with reproductive compensation and/or sibling competition (51), and more broadly males are thought to be “cursed” by mitochondrial alleles that can harm male fitness at a benefit to female fitness (52, 53). Similarly, natural selection also favors maternal effect dominant embryonic arrest alleles (MEDEA) that epigenetically program the arrest of embryos that do not inherit this allele (54). These scenarios are somewhat similar to the adaptive load model presented above, in that these alleles induce self harm based on contextual information of their situation to increase their inclusive fitness. However, contrary to the case of the adaptive load modelled here, both cytoplasmic male killers and MEDEA alleles increase their inclusive fitness at the expense of unlinked loci – setting up a genetic conflict between these selfish alleles and the rest of the genome. In our model of the adaptive load, the self-sacrificial allele increases the inclusive fitness of alleles at unlinked loci (at least for the maternally-derived genome). As such, rather than representing a genomic conflict, ours is a model of cooperation, and is consistent with Hamilton’s (55) idea that “infant mortality may evolve when the early death of one infant makes more likely the creation or survival of a close relative” (see also (24)). A closer analogue to our model is Charlesworth’s (26) model for the adaptive evolution of recessive lethal alleles linked to a driving t-haplotype, which would otherwise lead to sterility when homozygous.

Our results are reminiscent of models for the evolution of organismal robustness and sensitivity to deleterious mutations under differing forms of selection (56–58). For example, Archetti (57) showed that soft selection within families (akin to reproductive compensation in our model) favors alleles hypersensitive to deleterious mutations, and Lan et al (58) showed that asexual lineages that resist Mueller’s Ratchet by amplifying the fitness impacts of additional mutations by displaying synergistic epistasis can be favored by selection. However, in our model, alleles displaying sensitivity to homozygosity rather than mutation are favored.

### The efficacy of an altruistic load in preventing inbreeding

An adaptive recessive load can effectively prevent self-fertilization. For example, with a selfing rate of ten percent, lethal inbreeding, and complete recessivity and lethality of self-sacrificial alleles, we found about five such alleles per individual. Assuming these loci are unlinked, we expect an only 0.3125% of selfed seed to give rise to embryos for this parameter combination. However, the location, size, and selection coefficient of the mutational target for loci potentially underlying such early-acting inbreeding depression are unknown, so we have no firm predictions.

### Implications for the evolution of Self Incompatibility

Although we model the altruistic evolution of an early-expressed genetic load (rather than the spiteful load represented by cytoplasmic male killing and MEDEA), this evolutionary model is not without conflict. Here, the altruistic load clearly provides a greater patrilineal than matrilineal fitness cost, because an incidentally sacrificed embryo can be replaced by another from the same mother, while this pollen grain is replaced by one from a randomly chosen father. Although not directly modelled here, this conflict between maternally and paternally derived genes could favor the differential expression of the early embryonic load (i.e. genomic imprinting in embryos) depending on parent of origin (especially once the deleterious mutation is at high frequency), much like conflict over maternal resource provisioning favors parent-of-origin expression under the kinship theory of genomic imprinting (59). This view of the early adaptive load as a conflict between parental genomes over a self-sacrificial act provides a potential connection between self-incompatibility and the early expressed recessive load. This view is quite similar to Uyenoyama’s models that show that the differing costs and benefits of self-incompatibility alleles to pollen and style can favor pollen with novel SI specificities, (60, 61), consistent with the idea that pollen benefit more from having potentially low-fitness offspring than do mothers. This tension between pollen (or pollen parent) and style (or seed parent) over fertilization and development is not unique to self-fertilization – for example upon secondary contact the benefit to pollen in overcoming stylar incompatibilities and siring potentially low-fitness hybrids can frustrate the evolution of reinforcement (62).

### Testable predictions

> Adaptation is a special and onerous concept that should not be used unnecessarily.
>
> — George C. Williams. 1966.

#### *Adaptation and Natural Selection* (63)

Our model predicts that inadvertent selfing in predominantly outcrossing species with reproductive compensation can favor the adaptive evolution of early expressed recessive deleterious variation. However, our genome-scale simulations point to a more parsimonious explanation for an early expressed load. That is, when mutation rates are sufficiently high, the early expressed load that evolves by standard mutation-selection balance can equal or exceed the load favored by the adaptive process presented here, obviating the need for an adaptive load. Given that a recessive load can be efficiently generated by the process of mutation rather than selection, we urge for careful tests of the ideas developed here before accepting the adaptive load hypothesis. We present some testable predictions below.

#### We expect an elevated recessive embryonic load in taxa with effective reproductive compensation

The benefit of early embryonic death is that this death can be compensated by redirected resources that would go to this embryo to a current or future sibling. As such, we predict the load of early expressed recessive deleterious variants to increase in taxa with more reproductive compensation. However, while this observation is consistent with our adaptive load model, it is also consistent with the more parsimonious explanation that reproductive compensation relaxes selection against an early-expressed load (25, 27), counteracting the purging process in inbred populations (44).

#### Early-acting inbreeding depression should increase with the primary selfing rate (up to a point)

The greater the risk of inadvertent selfing, the greater the benefit of adaptive early-acting inbreeding depression. As such, we predict a larger early expressed recessive load in species with a higher selfing risk than those that effectively prevent the risk of self pollination. In such comparisons, the post-embryonic load – which should be unaffected by the process modelled above – could serve as a useful control. We note however, that we assumed that inbreeding depression was extreme and could not evolve. This occurs when selfing is rare and recessive mutations are common (31), has been observed in self-incompatible populations (e.g. 64), and is likely true of most primarily outcrossing populations. Because we do not model the co-evolution of mating system and the early genetic load, our predictions are only valid for taxa in which selfed seeds have very low fitness.

#### Recessive loci underlying late-acting inbreeding depression should show signatures of balancing selection

When the recessive load evolves adaptively (rather than under mutation selection balance) we see that recessive alleles are maintained under balancing selection and are on average older than those maintained under mutation selection balance. Thus, if the early expressed load is an adaptation, we predict a signature of balancing selection around loci underlying it – i.e. under a model of mutation-selection balance, recessive deleterious alleles should be relatively young, while under a model of an altruistic load, such alleles should be relatively old.

### Examination of the embryo lethality in gymnosperms

Our model is of particular relevance to gymnosperms, as they lack a self-incompatibility mechanism, cannot avoid geitonogamous selfing and display high inbreeding depression. Pines have one to four archegonia per ovule – of which all can be fertilized, with only one surviving to be included in the seed (65). Our model makes sense of Williams’ (66) conclusion that embryo death in pines occurs by two processes: (i) Numerous “Embryo Viability Loci” which are expressed throughout development and which potentially allow for embryo replacement, and (ii) The Embryo Lethal System – a coordinated process of embryo death which occurs after a dominant embryo has been established with limited opportunity for replacement of a dead embryo. We argue that polyembryony initially evolves as a mechanism to compensate the loss of embryos to inbreeding (as we 67, found previosuly), and that many of the “embryo viability loci” seen in nature represent the adaptive recessive load discovered in this work. Finally, we argue that the “embryo lethal system” is an evolved mechanism to suppress the non-dominant embryo if both embryos are viable – an interpretation consistent with both our theoretical observation that embryo competition frustrates the evolution of a recessive load (67), and the claim that death induced by the embryo lethal system is regulated by the position of the embryo rather than its competitive ability (66). This interpretation of a recessive load as an inbreeding avoidance strategy, is also consistent with the observation that gymnosperms have a lower realized selfing rate than expected given their primary selfing rates (68), and an intense study of a pine orchard demonstrating that the recessive load, rather than the opportunity for self-fertilization, controlled the realized selfing rate (69)).

### What proportion of the early-expressed load is altruistic?

We have show that an altruistic,”self-sacrificial,” recessive genetic load *can* evolve by natural selection. We again reiterate that this does not mean that much of the early-expressed load is adaptive. The proportion of the load which is self-sacrificial likely varies tremendously across taxa and depends on the extent of reproductive compensation (more compensation should yield a more altruistic load), the mutation rate to deleterious mutations that allow for compensation (the lower this rate, the more of the load can be attributable to selection rather than mutation), and the extent of inbreeding and inbreeding depression.

## Summary

We show that when mothers can compensate for inviable embryos, a self-sacrificial early expressed recessive load can adaptively evolve by natural selection. Our model shows that the line between “early-acting inbreeding depression” and “late-acting self-incompatibility” might not be as conceptually clear-cut as previous authors have insisted. As such, our model potentially refutes the orthodox idea that “Abortion of inbred embryos is not an evolved trait for ensuring outcrossing, but is considered an early expression of genetic load, a lethal ‘price’ a species pays for maintaining genetic variability” (22) – and the price paid is the that of incomplete reproductive compensation. However, our model also shows that such a mechanism only meaningfully operates when the rate of origin of such mutations is low, as selection does not favor such mutations when they arise so frequently that a substantial recessive load would evolve in the absence of selection. We therefore argue that the key evolutionary question is not to distinguish between early-acting inbreeding depression and late acting self-incompatibility, but rather to distinguish between adaptive and non-adaptive mechanisms that prevent the production of selfed seed.

## ACKNOWLEDGEMENTS

We would like to acknowledge Boris Igic, Mike Wade, Husain Agha, Aidan Harrington, Brooke Kern, and Manisha Munasinghe, and other members of the Brandvain lab for comments and the Research Council of Finland for supporting our project ‘Genetic determinants of embryo destiny in a polyembryonic seed – mortality, inbreeding depression and sibling rivalry in a pine nutshell’ (349057). Previous drafts are available on bioRxiv (70), and code underlying the results are available on zenodo (30).

## Appendix

This mathematical appendix accompanies the manuscript, Brandvain, Y., Thomson, L., and Pyhäjärvi, T. 2024. Early-acting inbreeding depression can evolve as an inbreeding avoidance mechanism.

### Mating Table, Single locus model

Here we show all possible offspring genotypes, their adult and embryo fitness, and their frequency in each maternally family after early embryo selection. Maternal families occur in frequencies of their genotypes after selection. The relative fitness of a selfed plant equals 1 *− δ*, which we assume to equal zero throughout the analytical portion of the manuscript.

The values, *ψ*_*aa*_, *ψ*_*Aa*_, and *ψ*_*AA*_ denote the probability that an embryo is replaced by another (i.e. compensated) in *aa, Aa*, and *AA* families, respectively. The value, *ψ* in maternal family, *k* equals replace*k* =Σ Σfreq_Geno:i, of type: j, in Fam:k_ *×* embryo fitness*i* summed across all primary embryos for each maternal family. These values are: *ψ*_*aa*_ = *phs*(1 *− σ*), *ψ*_*Aa*_ = *s*(2*h* + 2*p*(1 *− σ*) + *σ*)*/*4, and *ψ*_*AA*_ = *s*(*p* + (1 *− p*)(*h*(1 *− σ*) + *σ*)).

| maternal fam | genotype | type | adult fitness | primary | embryo fitness | freq in fam |
| --- | --- | --- | --- | --- | --- | --- |
| aa | aa | Self | $1 - \delta$ | Primary | 1 | $\sigma$ |
| aa | Aa | Self | $1 - \delta$ | Primary | $1 - hs$ | 0 |
| aa | AA | Self | $1 - \delta$ | Primary | $1 - s$ | 0 |
| aa | aa | Out | 1 | Primary | 1 | $(1 - p)(1 - \sigma)$ |
| aa | Aa | Out | 1 | Primary | $1 - hs$ | $p(1 - \sigma)$ |
| aa | AA | Out | 1 | Primary | $1 - s$ | 0 |
| aa | aa | Self | $1 - \delta$ | Backup | 1 | $\psi_{aa}\sigma$ |
| aa | Aa | Self | $1 - \delta$ | Backup | $1 - hs$ | 0 |
| aa | AA | Self | $1 - \delta$ | Backup | $1 - s$ | 0 |
| aa | aa | Out | 1 | Backup | 1 | $\psi_{aa}(1 - \sigma)(1 - p)$ |
| aa | Aa | Out | 1 | Backup | $1 - hs$ | $\psi_{aa}(1 - \sigma)p$ |
| aa | AA | Out | 1 | Backup | $1 - s$ | 0 |
| Aa | aa | Self | $1 - \delta$ | Primary | 1 | $\sigma/4$ |
| Aa | Aa | Self | $1 - \delta$ | Primary | $1 - hs$ | $\sigma/2$ |
| Aa | AA | Self | $1 - \delta$ | Primary | $1 - s$ | $\sigma/4$ |
| Aa | aa | Out | 1 | Primary | 1 | $(1 - \sigma)(1 - p)/2$ |
| Aa | Aa | Out | 1 | Primary | $1 - hs$ | $(1 - \sigma)/2$ |
| Aa | AA | Out | 1 | Primary | $1 - s$ | $(1 - \sigma)(p/2)$ |
| Aa | aa | Self | $1 - \delta$ | Backup | 1 | $\psi_{Aa}(\sigma/4)$ |
| Aa | Aa | Self | $1 - \delta$ | Backup | $1 - hs$ | $\psi_{Aa}(\sigma/2)$ |
| Aa | AA | Self | $1 - \delta$ | Backup | $1 - s$ | $\psi_{Aa}(\sigma/4)$ |
| Aa | aa | Out | 1 | Backup | 1 | $\psi_{Aa}(1 - \sigma)(1 - p)/2$ |
| Aa | Aa | Out | 1 | Backup | $1 - hs$ | $\psi_{Aa}(1 - \sigma)/2$ |
| Aa | AA | Out | 1 | Backup | $1 - s$ | $\psi_{Aa}(1 - \sigma)p/2$ |
| AA | aa | Self | $1 - \delta$ | Primary | 1 | 0 |
| AA | Aa | Self | $1 - \delta$ | Primary | $1 - hs$ | 0 |
| AA | AA | Self | $1 - \delta$ | Primary | $1 - s$ | $\sigma$ |
| AA | aa | Out | 1 | Primary | 1 | 0 |
| AA | Aa | Out | 1 | Primary | $1 - hs$ | $(1 - p)(1 - \sigma)$ |
| AA | AA | Out | 1 | Primary | $1 - s$ | $p(1 - \sigma)$ |
| AA | aa | Self | $1 - \delta$ | Backup | 1 | 0 |
| AA | Aa | Self | $1 - \delta$ | Backup | $1 - hs$ | 0 |
| AA | AA | Self | $1 - \delta$ | Backup | $1 - s$ | $\psi_{AA}\sigma$ |
| AA | aa | Out | 1 | Backup | 1 | 0 |
| AA | Aa | Out | 1 | Backup | $1 - hs$ | $\psi_{AA}(1 - \sigma)(1 - p)$ |
| AA | AA | Out | 1 | Backup | $1 - s$ | $\psi_{AA}(1 - \sigma)p$ |

### Selection, Single locus model

The frequency of each genotype after selection is the product of its frequency in a maternal family, the maternal family’s frequency, its embryonic and adult fitness, weighted by mean fitness. By appropriately summing and weighting the content of the table above, and assuming complete inbreeding depression (*δ* = 1), we find mean fitness equals

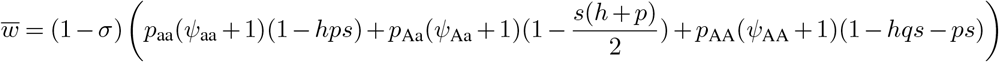

Where *σ* is the selfing rate, *p* and *q* are frequencies of alleles *a* (wild type), and *A* (self-sacrificial), respectively, and *ψ*_*k*_ is the probability that a primary embryo in the family of the subscripted maternal genotype, _*k*_ is replaced.

From the table above (see algebraic details in our Supplemental Mathematica file), we find that the frequencies of each genotype in the next generation are as follows:

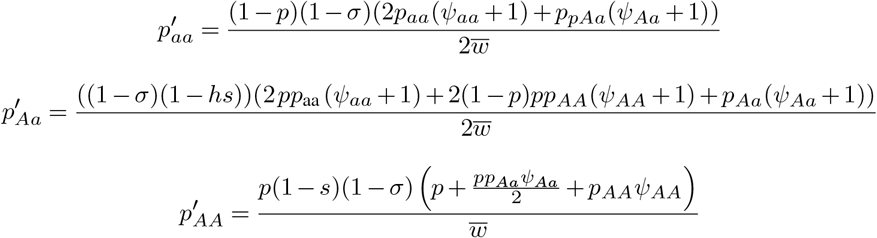

## Supplementary figures

These supplementary figures accompany the manuscript, Brandvain, Y., Thomson, L., and Pyhäjärvi, T. 2024. Early-acting inbreeding depression can evolve as an inbreeding avoidance mechanism.

**Fig. S1:**
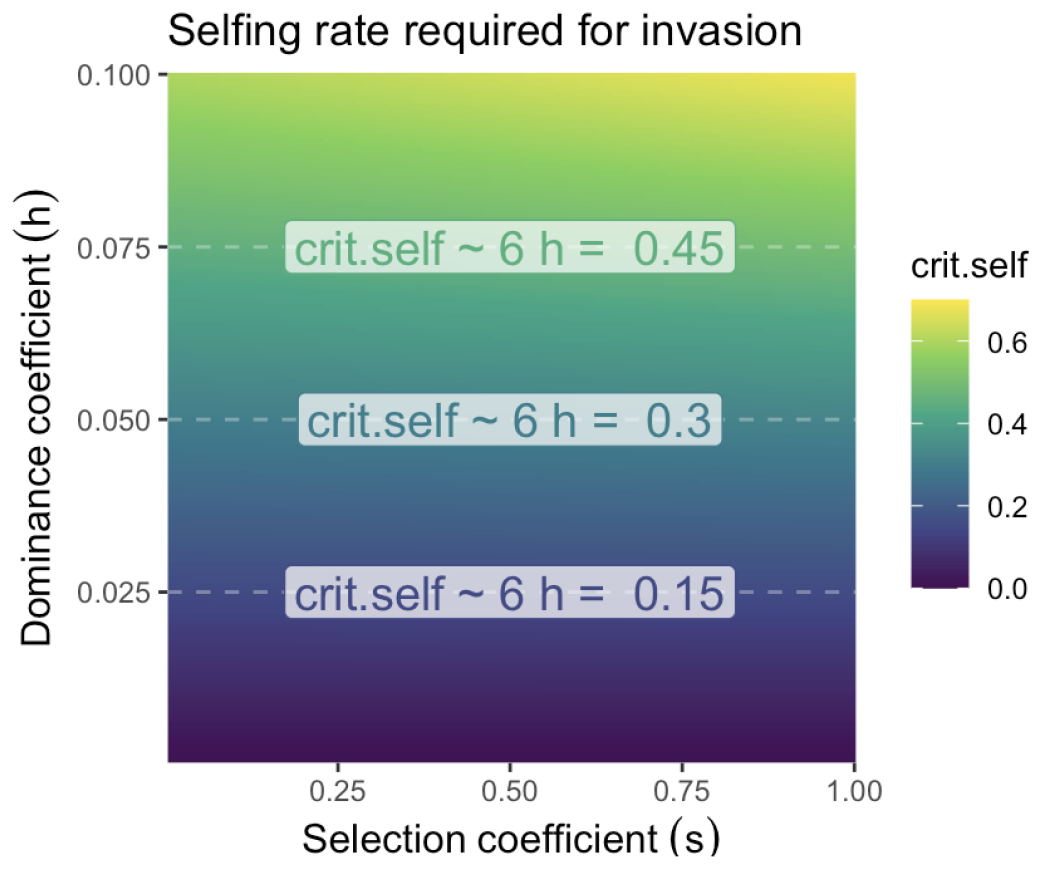
The critical selfing threshold required for the invasion of a self-sacrificial, *A*, allele as a function of its selection and dominance coefficients, under our model and assuming that all selfed seed dies. The colored text reflects the approximation that a rare *A* allele is favored when crit.self *>* 6*h*. This approximation is exact as *s* approaches zero, but crit.self must exceed 6*h* as *s* increases (e.g. with *s* = 1 crit.self equals 0.155, 0.321, and 0.499, when *h* is 0.025, 0.05, and 0.075, respectively.)

**Fig. S2:**
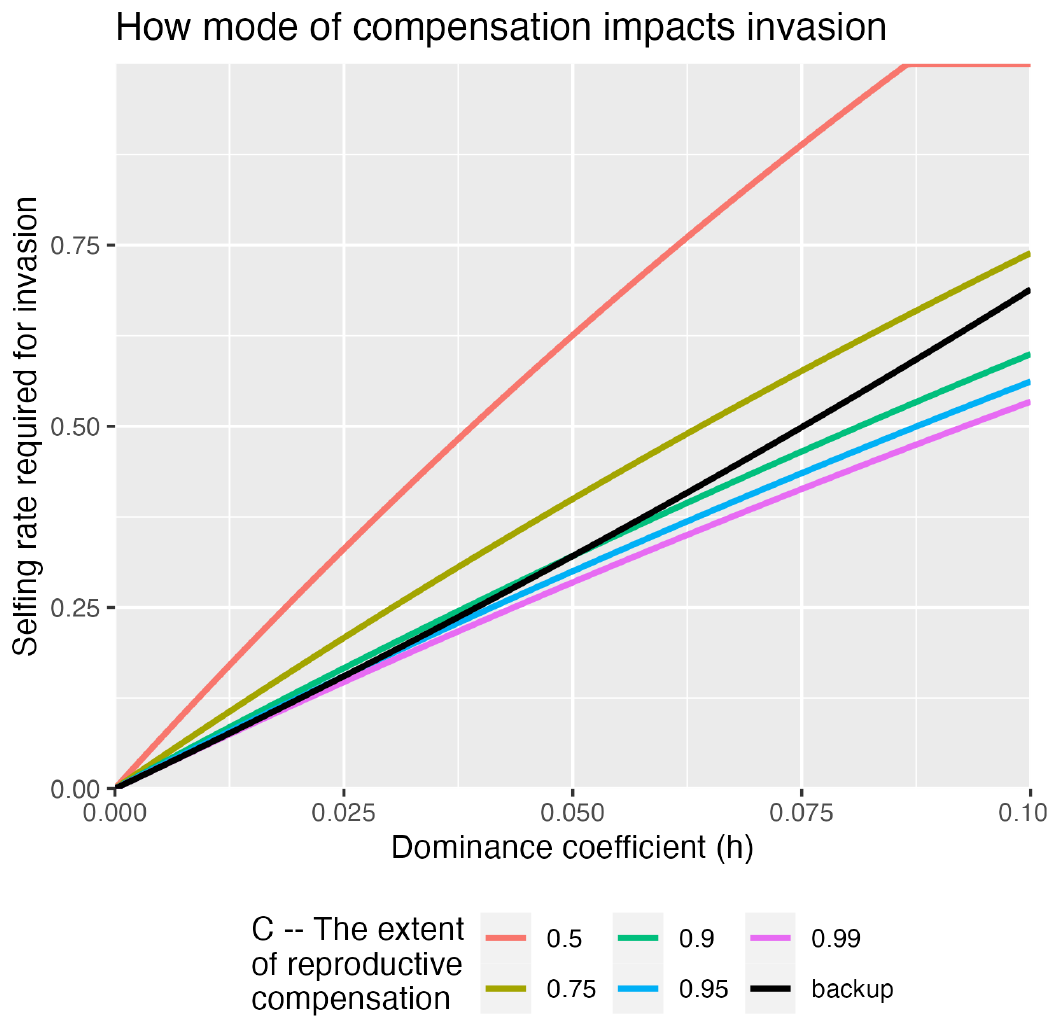
How the extent and mechanism of reproductive compensation impact the invasion criteria of a self-sacrificial (partially) recessive allele across dominance coefficients. The level of compensation, *C*, in Charlesworth’s parameterization of reproductive compensation is shown in color, while results from our parameterization of reproductive compensation are provided in black as a reference. Results – reported as the critical selfing rate required for invasion (y-axis) are shown for a variety of dominance coefficients (x-axis). All lines assume a selection coefficient of *s* = 0.5.

**Fig. S3:**
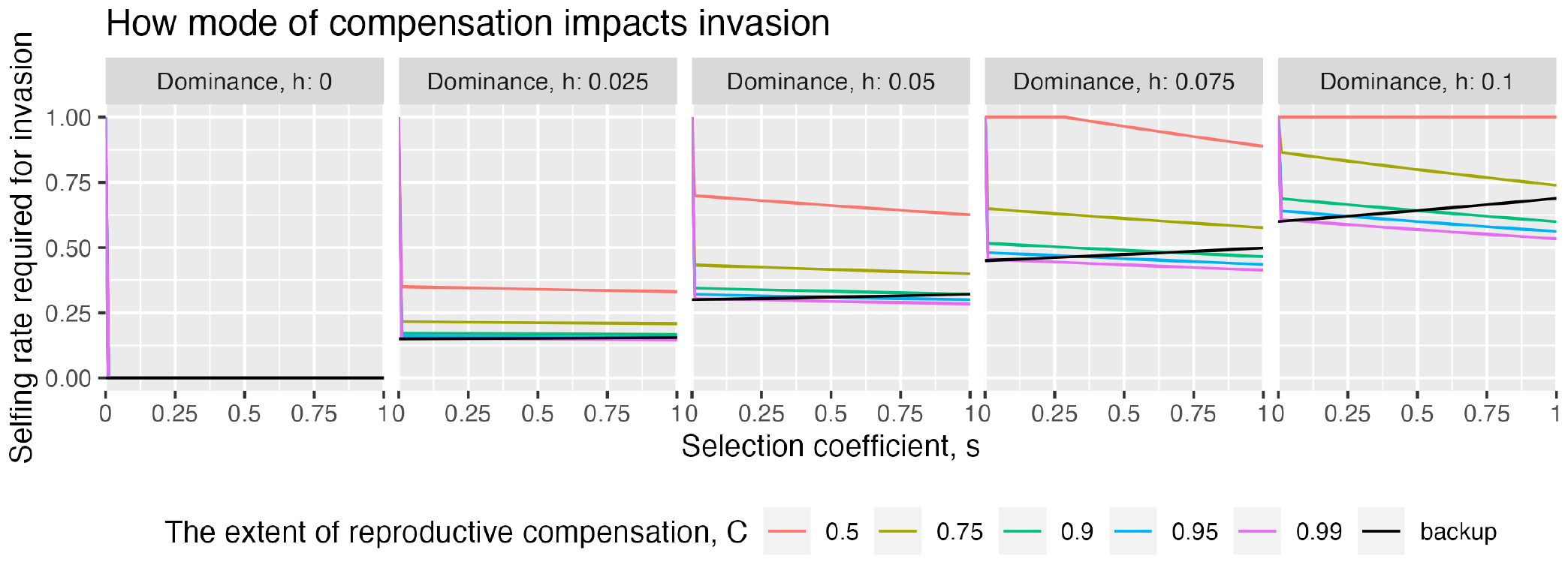
How the extent and mechanism of reproductive compensation impact the invasion criteria of a self-sacrificial (partially) recessive allele across selection and dominance coefficients. The level of compensation, *C*, in Charlesworth’s parameterization of reproductive compensation is shown in color is show in color, while results from our parameterization of reproductive compensation are provided in black as a reference. Results – reported as the critical selfing rate required for invasion (y-axis) are shown for a variety of selection coefficients (x-axis) and dominance coefficients (facets).

**Fig. S4:**
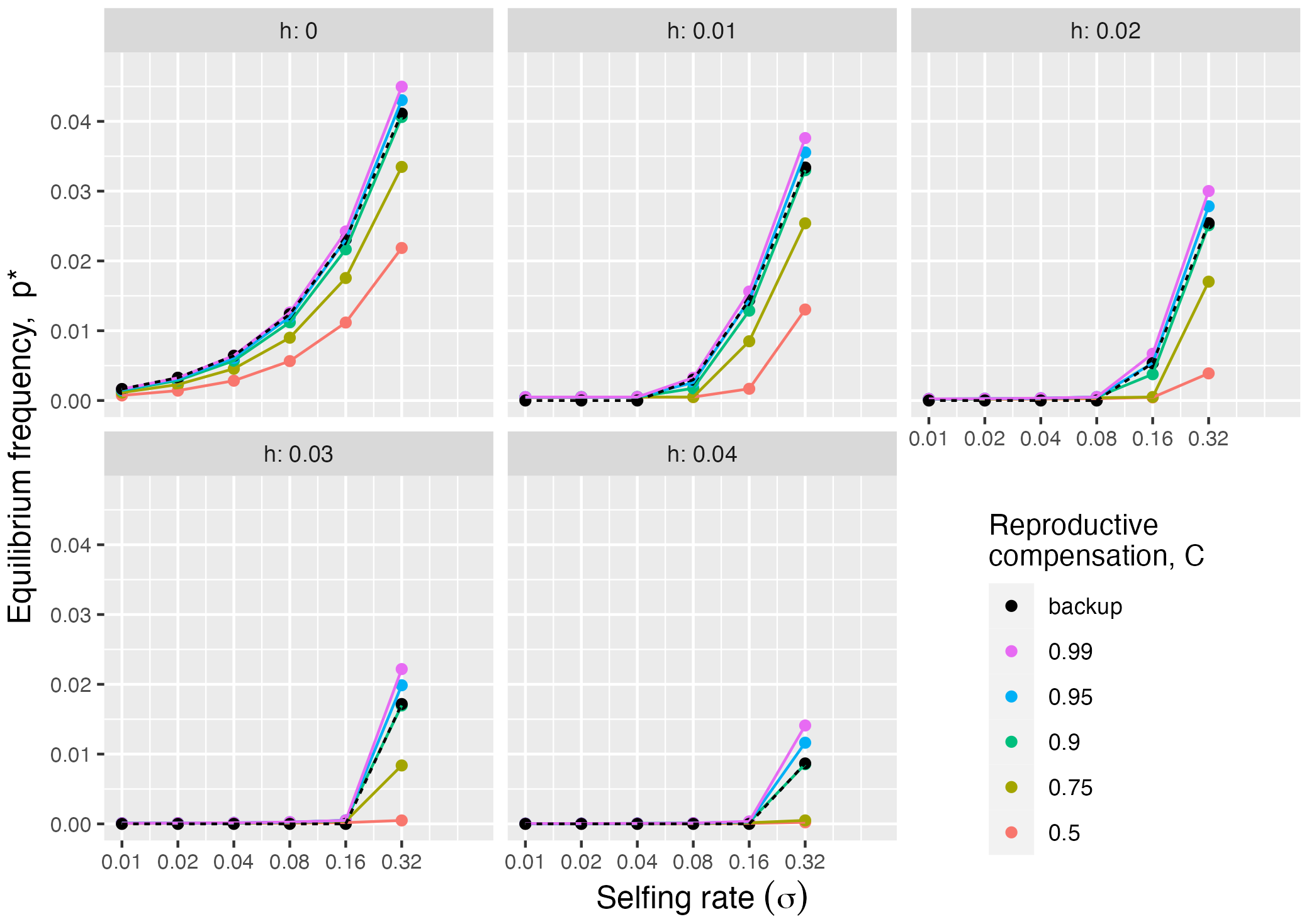
How the extent and mechanism of reproductive compensation impact the equilibrium frequency, *p*^*∗*^, of a self-sacrificial (partially) recessive lethal allele. The level of compensation, *C*, in Charlesworth’s parameterization of reproductive compensation is shown in color is show in color, while results from our parameterization of reproductive compensation are provided in black as a reference. Results are shown for a variety of selfing rates (x-axis) and dominance coefficients (facets), and assume lethal inbreeding.

**Fig. S5:**
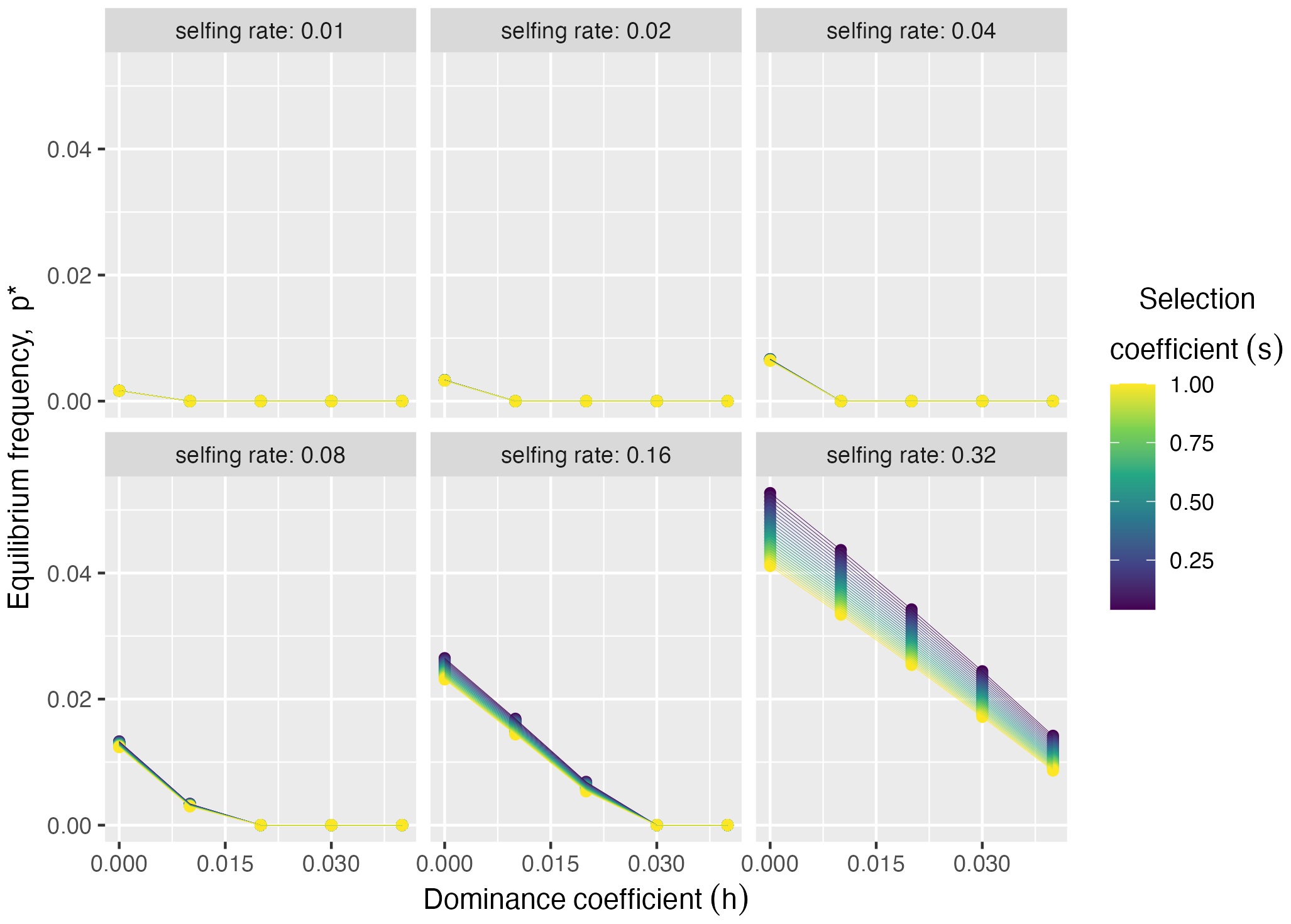
The equilibrium frequency, *p*^*∗*^, of a self-sacrificial (partially) recessive allele is largely insensitive to the strength of selection (in color), and depends more on the selfing rate (facet) and dominance coefficient (x-axis).

**Fig. S6:**
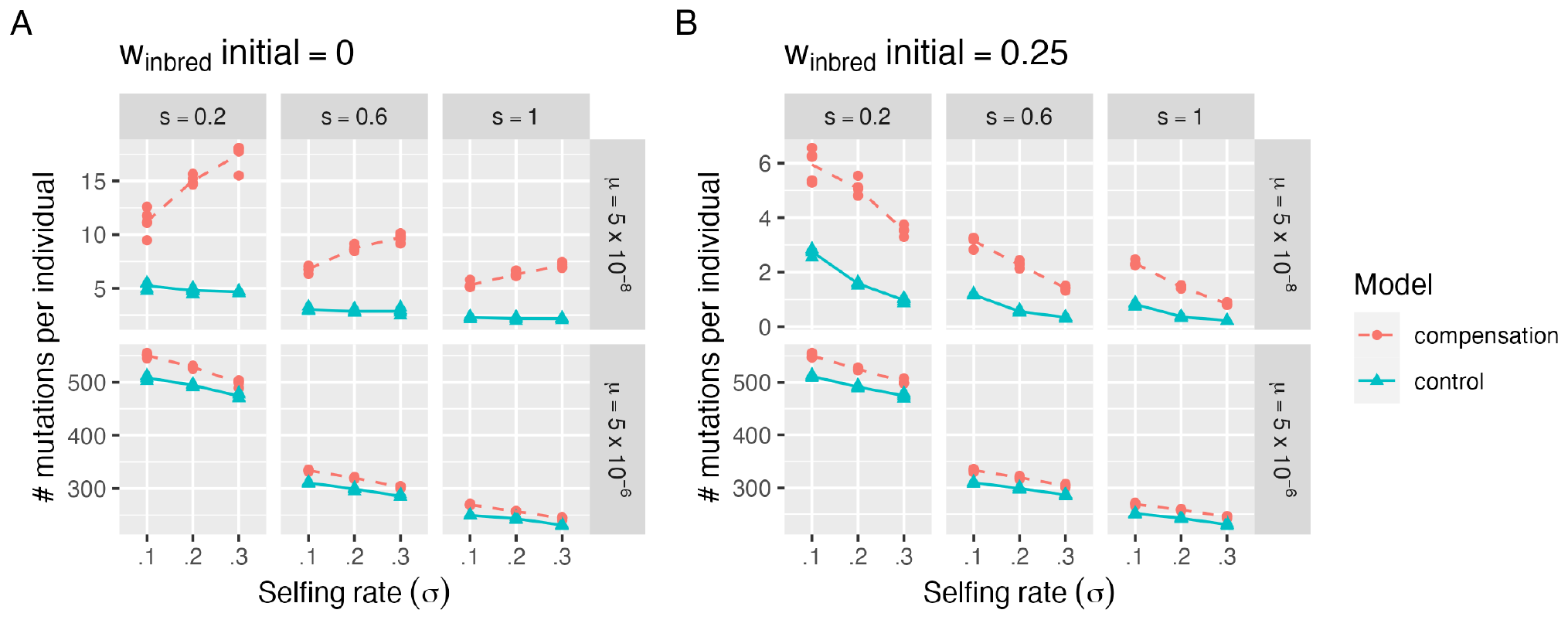
The mean number of early-acting, recessive lethal (*w*_inbred_ initial = 0, A), and sublethal (*w*_inbred_ initial = 0.25, B) alleles per individual, over a combination of selfing rates (*σ*), mutation rates (*µ*), selection (*s*)and dominance (h) coefficients. Blue lines display results for cases in which primary embryos are replaced by siblings (our model), while blue lines display cases in which primary embryos are replaced by a non-sibling (a control). Simulations were run in a population of size 10000.

**Fig. S7:**
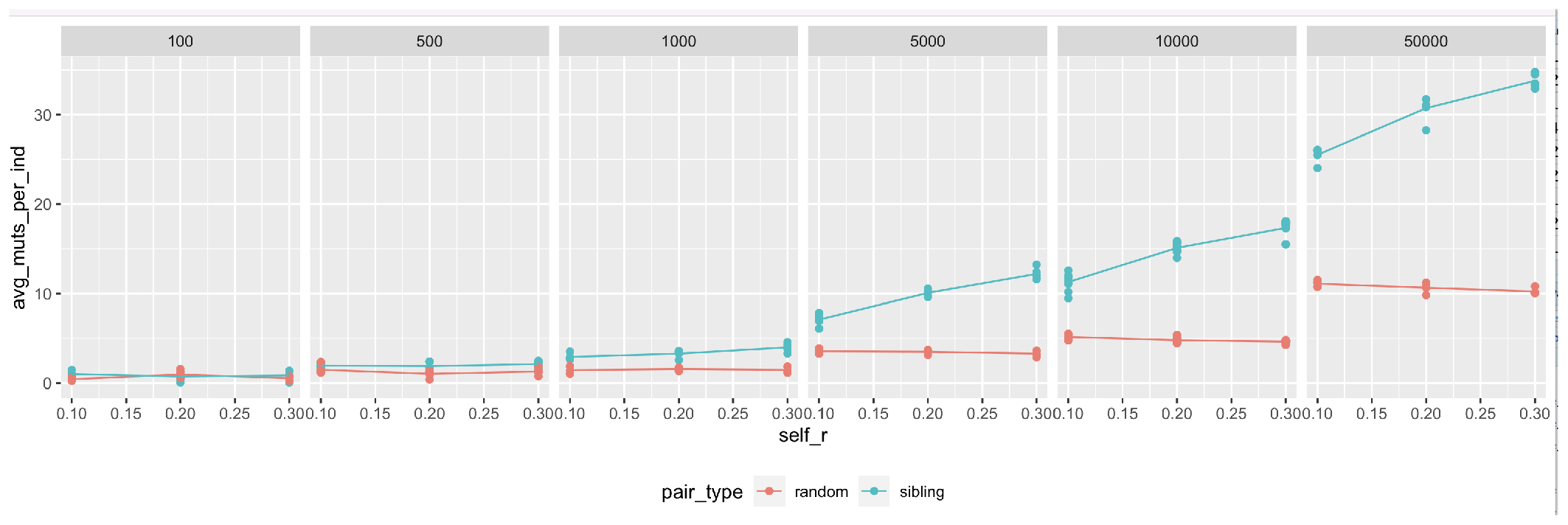
The mean frequency of early-acting, recessive alleles over a combination of selfing rates (*σ*, x-axis), and population sizes (facets). Blue lines display results for cases in which primary embryos are replaced by siblings (our model), while blue lines display cases in which primary embryos are replaced by a non-sibling (a control). Simulations assume lethal inbreeding and a selection coefficient of -0.2 on the recessive allele.

**Fig. S8:**
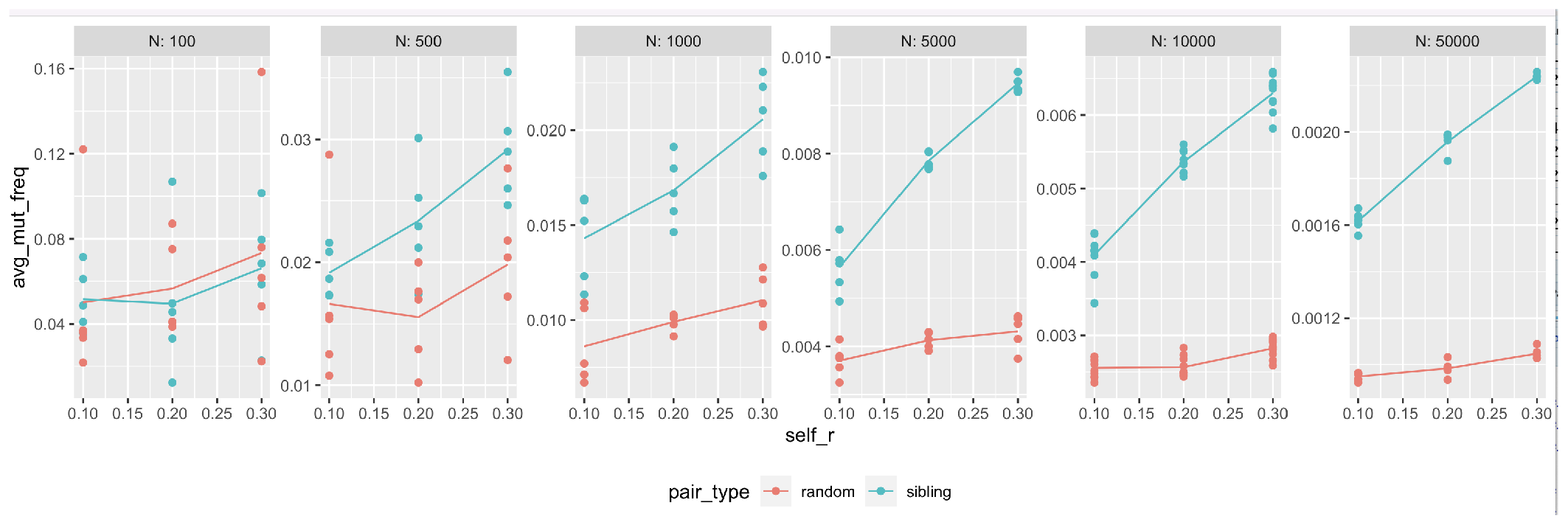
The mean number recessive alleles per individual over a combination of selfing rates (*σ*, x-axis), and population sizes (facets). Blue lines display results for cases in which primary embryos are replaced by siblings (our model), while blue lines display cases in which primary embryos are replaced by a non-sibling (a control). Simulations assume lethal inbreeding and a selection coefficient of -0.2 on the recessive allele.

## Supplementary text

This text accompanies the manuscript, Brandvain, Y., Thomson, L., and Pyhäjärvi, T. 2024. Early-acting inbreeding depression can evolve as an inbreeding avoidance mechanism.

### An alternative parameterization of reproductive compensation

To evaluate the robustness of our results to our model of reproductive compensation, we developed a second model with an alternative form of reproductive compensation. Specifically we implemented reproductive compensation following Charlesworth’s (26), model, in which maternal fitness equals 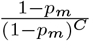, where *p*_*m*_ is the probability that an embryo will not survive and *C* is the extent of reproductive compensation (Fig Supp Text 1). Each mother produces *σ* of its children by selfing and 1 *− σ* of its children by outcrossing (receiving pollen at random). Each mother produces (up to) six type of embryos – selfed or outcrossed embryos of all three genotypes – in frequencies given in Table 1:

**Table 1.**
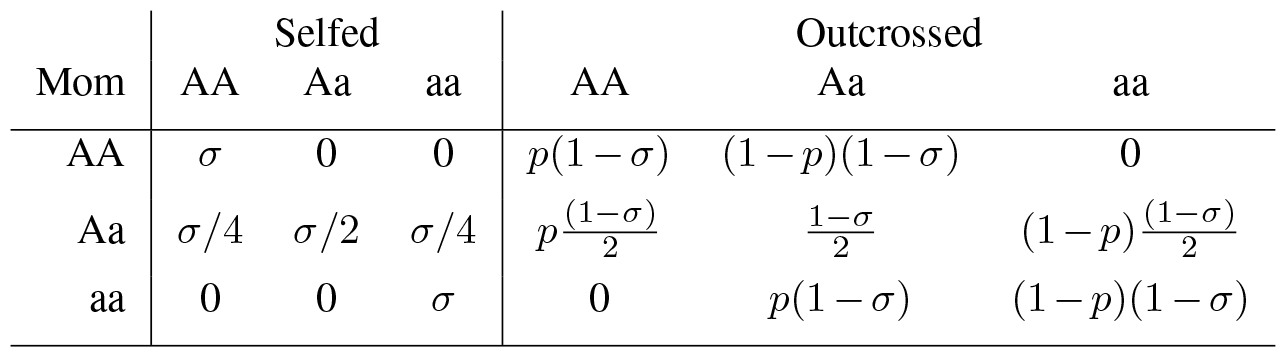
Frequencies of types of embryos in each maternal family before selection within families.

| Mom | Selfed |  |  | Outcrossed |  |  |
| --- | --- | --- | --- | --- | --- | --- |
|  | AA | Aa | aa | AA | Aa | aa |
| AA | $\sigma$ | 0 | 0 | $p(1-\sigma)$ | $(1-p)(1-\sigma)$ | 0 |
| Aa | $\sigma/4$ | $\sigma/2$ | $\sigma/4$ | $p\frac{(1-\sigma)}{2}$ | $\frac{1-\sigma}{2}$ | $(1-p)\frac{(1-\sigma)}{2}$ |
| aa | 0 | 0 | $\sigma$ | 0 | $p(1-\sigma)$ | $(1-p)(1-\sigma)$ |

We then select within families, which in our model depends only on genotype at the *𝒜* locus. Frequencies within families after selection equal frequencies before selection multiplied by the fitness of each genotype (during early development - i.e. depending on genotype, not selfed status) divided by mean family fitness. Recall that genotypic fitnesses during within family selection are (1 *− s*), (1 *− hs*), and 1 for AA, Aa, and aa genotypes, respectively. The denominators (i.e. mean family fitnesses before accounting for compensation) are therefore

After finding the relative frequencies of each combination of selfing and genotype in each family after selection, we introduce reproductive compensation following the model of Charlesworth (26). Here, the fitness of each maternal family *j* after compensation, *w*_Fam:j,AC_, becomes its value before compensation, *w*_Fam:j,AC_ divided by this value raised to the *C* (Eq. 1).

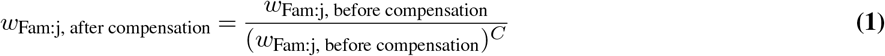

where *C* is a measure of the degree of compensation (if *C* = 1, early mortality has no effect on final offspring number, while if *C* = 0 there is no reproductive compensation).

We then find the frequency of each of the six offspring types after selection within and among families (i.e. in dispersed seed in our model) as the product of its frequency within a family after within family selection multiplied by the contribution of this family to the seed pool (i.e. after among family after selection, where this frequency is simply the family frequency before selection multiplied by its fitness after compensation divided by the mean fitness of all families after compensation), summed across all families. So, the frequency of offspring type, *i*, after selection within and among families equals

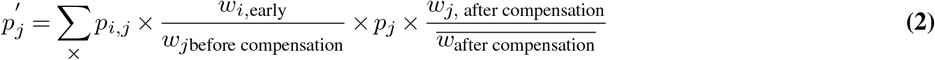

where *p*_*i,j*_ is the frequency of offspring type, *i* in family *j* before selection (i.e. values in Table 2), *w*_*i*,early_ is the fitness of this offspring (during within family selection), based on its genotype, *w*_*j*before compensation_ corresponds to family fitness values before compensation presented in Table 2, while *w*_*j*, after compensation_ refers to values derived from Eq. 1, and 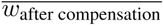 is the mean of Eq. 2 (i.e. this value for each family multiplied by the family frequency before selection, summed across all families).

**Table 2.** Mean fitness of each family before reproductive compensation.

| Mom | mean fitness |
| --- | --- |
| AA | $1 - s(\sigma + (1-\sigma)(p + h(1-p)))$ |
| Aa | $1 - \frac{s}{2}(h + p(1-\sigma + \sigma/2))$ |
| aa | $1 - hps(1-\sigma)$ |

**Fig Supp Text 1:**
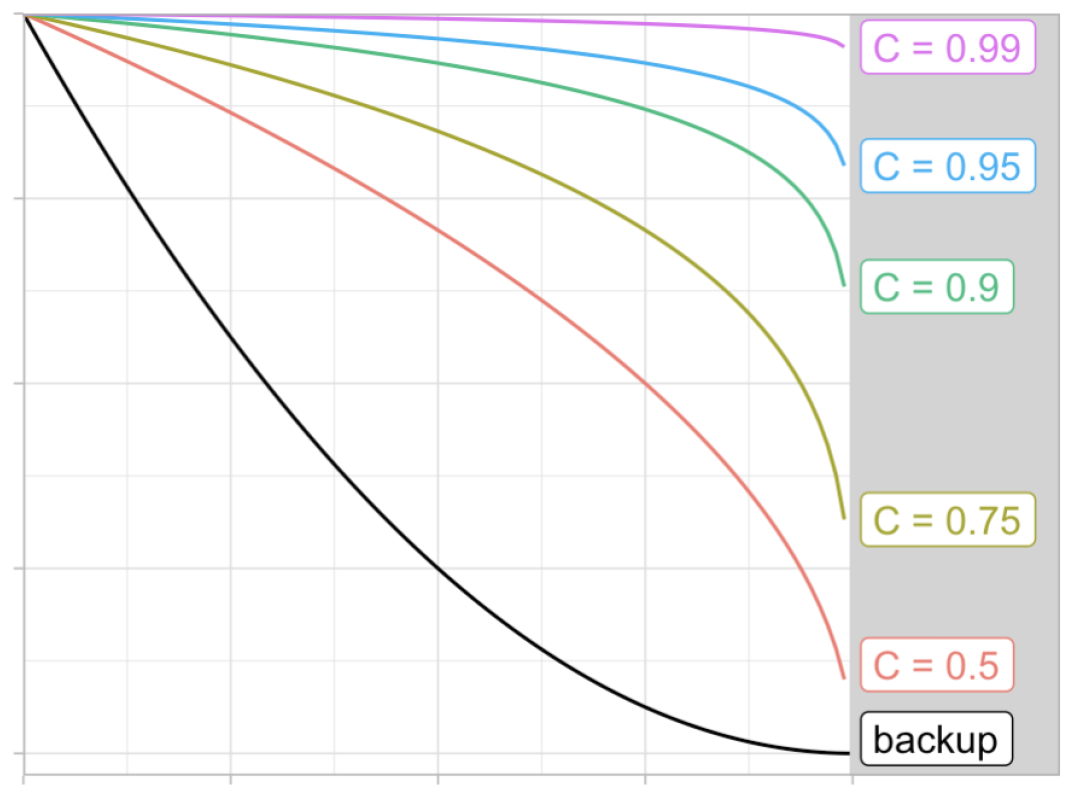
The extent of reproductive compensation as a function of *C* (color) under Charlesworth’s parameterization of reproductive compensation, in which maternal fitness equals (1 *− p*_*m*_)*/*((1 *− p*_*m*_)^*C*^).

Finally, surviving seeds are subjected to later selection, which we base exclusively on their inbreeding status. So the offspring types after this stage of selection, 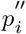 equal 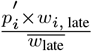. Because this math is quite cumbersome, we find results for invasion and equilibrium allele frequencies by numerical iteration in R.

### Invasion analysis for Charlesworth’s parameterization of reproductive compensation

Because this parameterization model was not analytically tractable, we used a bisection optimization approach to identify invasion criteria. For each combination of parameters, we found critical selfing rate required for invasion as follows:

1. We kept track of three values – our focal selfing rate (initially 0.5), our lower selfing rate (initially 0), and our upper selfing rate (initially 1).
2. We evaluated invasion by introducing (at frequency .000005) a deleterious early expressed allele, and asked if it deterministically increased in frequency from generation one to ten given the focal selfing rate. And then
  - If the allele invaded, we updated our values as follows:
    -*The focal selfing rate* takes a value halfway between the previous focal selfing rate and the lower selfing rate.
    -*The upper selfing rate* takes the value of the previous focal selfing rate.
    -*The lower selfing rate* does not change.
  - If the allele failed to invade, we updated our values as follows:
    -*The focal selfing rate* takes a value halfway between the previous focal selfing rate and the upper selfing rate.
    -*The upper selfing rate* does not change.
    -*The lower selfing rate* takes the value of the previous focal selfing rate.
3. After a test at one focal selfing rate we then evaluate invasion at the new focal selfing rate. We continue this process until the difference between the upper and lower selfing rates is less than 10 ^0*−*4^ – a precision usually achieved in about fifteen iterations. We consider the focal selfing rate at this point to be the lowest selfing rate which allows for invasion (i.e. the “critical selfing rate”).

## Notes

https://zenodo.org/records/10552087

### Competing Interest Statement

The authors have declared no competing interest.

### Summary of Updates

We showed that the efficacy o selection for an altrusitic load is most effective in large populations.

